# Comparative assessment of genes driving cancer and somatic evolution in noncancer tissues: an update of the NCG resource

**DOI:** 10.1101/2021.08.31.458177

**Authors:** Lisa Dressler, Michele Bortolomeazzi, Mohamed Reda Keddar, Hrvoje Misetic, Giulia Sartini, Amelia Acha-Sagredo, Lucia Montorsi, Neshika Wijewardhane, Dimitra Repana, Joel Nulsen, Jacki Goldman, Marc Pollit, Patrick Davis, Amy Strange, Karen Ambrose, Francesca D. Ciccarelli

## Abstract

Genetic alterations of somatic cells can drive nonmalignant clone formation and promote cancer initiation. However, the link between these processes remains unclear hampering our understanding of tissue homeostasis and cancer development. Here we collect a literature-based repertoire of 3355 well-known or predicted drivers of cancer and noncancer somatic evolution in 122 cancer types and 12 noncancer tissues. Mapping the alterations of these genes in 7953 pancancer samples reveals that, despite the large size, the known compendium of drivers is still incomplete and biased towards frequently occurring coding mutations. High overlap exists between drivers of cancer and noncancer somatic evolution, although significant differences emerge in their recurrence. We confirm and expand the unique properties of drivers and identify a core of evolutionarily conserved and essential genes whose germline variation is strongly counter-selected. Somatic alteration in even one of these genes is sufficient to drive clonal expansion but not malignant transformation. Our study offers a comprehensive overview of our current understanding of the genetic events initiating clone expansion and cancer revealing significant gaps and biases that still need to be addressed. The compendium of cancer and noncancer somatic drivers, their literature support and properties are accessible at http://www.network-cancer-genes.org/.

## BACKGROUND

Genetic alterations conferring selective advantages to cancer cells are the main drivers of cancer evolution and hunting for them has been at the core of international cancer genomic efforts ^1,2,3^. Given the instability of the cancer genome, distinguishing driver alterations from the rest relies on analytical approaches that identify genes altered more frequently than expected or quantify the positive selection acting on them ^4,5,6^. The results of these analyses have greatly expanded our understanding of the mechanisms driving cancer evolution, revealing high heterogeneity across and within cancers ^7,8,9^.

Recently, deep sequencing screens of noncancer tissues have started to map positively selected genetic mutations in somatic cells that drive *in situ* formation of phenotypically normal clones ^10,11^. Many of these mutations hit cancer drivers, sometimes at a frequency higher than in the corresponding cancer ^12,13,14,15,16^. Yet, they do not drive malignant transformation. This conundrum poses fundamental questions on how genetic drivers of normal somatic evolution are related to and differ from those of cancer evolution. Addressing these questions will clarify the genetic relationship between tissue homeostasis and cancer initiation, with profound implications for cancer early detection.

To assess the extent of the current knowledge on cancer and noncancer drivers, we undertook a systematic review of the literature and assembled a comprehensive repertoire of genes whose somatic alterations have been reported to drive cancer or noncancer evolution. This allowed us to compare the current driver repertoire across and within cancer and noncancer tissues and map their alterations in the large pancancer collection of samples from The Cancer Genome Atlas (TCGA). This revealed significant gaps and biases in our current knowledge of the driver landscape. We also computed an array of systems-level properties across driver groups, confirming the unique evolutionary path of driver genes and their central role in the cell.

We collected all cancer and noncancer driver genes, together with a large set of their properties, in the Network of Cancer Genes and Healthy Drivers (NCG^HD^) open-access resource.

## RESULTS

### More than 3300 genes are canonical or candidate drivers of cancer and noncancer somatic evolution

We conducted a census of currently known drivers through a comprehensive literature review of 331 scientific articles published between 2008 and 2020 describing somatically altered genes with a proven or predicted role in cancer or noncancer somatic evolution (**Figure 1A**). These publications included three sources of experimentally validated (canonical) cancer drivers, 311 sequencing screens of cancer (293) and noncancer (18) tissues and 17 pancancer studies (**Supplementary Table 1, Additional File 1**). Each paper was assessed by at least two independent experts (**Supplementary Figure 1A-C, Additional File 2**) returning a total of 3355 drivers, 3347 in 122 cancer types and 95 in 12 noncancer tissues, respectively (**Figure 1A**). We further computed the systems-level properties of drivers and annotated their function, somatic variation and drug interactions (**Figure 1A**).

**Figure 1.**
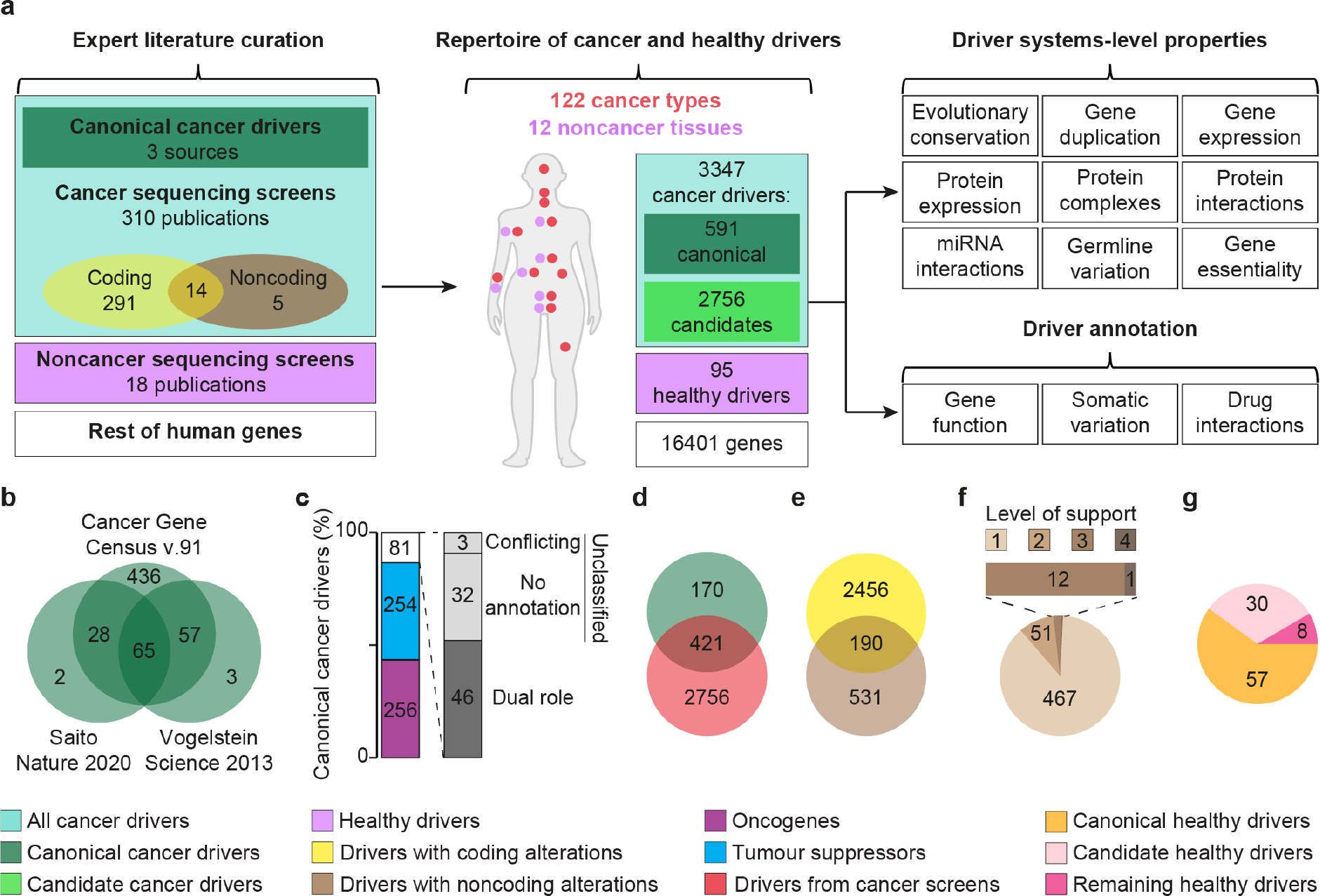
Collection of a comprehensive repertoire of cancer and healthy drivers. **a**. Literature review and driver annotation workflow. Expert literature curation of 331 publications led to a repertoire of cancer and healthy drivers in a variety of cancer and noncancer tissues. Combining multiple data sources, a set of properties and annotations were computed for all these drivers. **b**. Intersection of canonical drivers from three sources ^17,18,19^ that passed our manual curation. **c**. Classification of canonical cancer drivers in tumour suppressors and oncogenes. Eighty-one cancer drivers had a dual role or could not be classified. **d**. Intersection of canonical and candidate driver genes from 310 sequencing screens. Genes whose driver role had only statistical support were considered candidate cancer drivers. **e**. Intersection between cancer drivers with coding and noncoding alterations. **f**. Level of support for the driver role of 531 cancer genes with noncoding driver alterations only. Level 1 means that the gene was predicted as a driver only in one cancer sequencing screen; levels 2, 3 and 4 mean that it was predicted by two, three or four screens or that it had experimental support. Experimental support was gathered from the 19 publications reporting noncoding cancer drivers (**Supplementary Table 1, Additional File 1**) and form the CNCDatabase ^20^ and included in vitro and in vivo experiments, modification of gene expression and survival association. **g**. Proportion of healthy drivers that are also canonical or candidate cancer drivers, classified as canonical and candidate healthy drivers, respectively.

We reviewed the three sources of canonical cancer drivers ^17,18,19^ to exclude false positives (**Supplementary Table 2, Additional File 1**) and fusion genes whose properties could not be mapped. Only 11% of the resulting 591 canonical drivers (**Supplementary Table 3, Additional File 1**) were common to all three sources (**Figure 1B**), indicating poor consensus even in well-known cancer genes. We further annotated the genetic mode of action for >86% of canonical drivers, finding comparable proportions of oncogenes or tumour suppressors (**Figure 1C**). The rest had a dual role or could not be univocally classified.

We extracted additional cancer drivers from the curation of 310 sequencing screens that applied a variety of statistical approaches (**Supplementary Figure 1D, Additional File 2**) to identify cancer drivers among all altered genes. After removing possible false positives (**Supplementary Table 2, Additional File 1**), the final list included 3177 cancer drivers, 2756 of which relied only on statistical support (candidate cancer drivers) and 421 were canonical drivers (**Figure 1D, Supplementary Table 3, Additional File 1**). Therefore, 170 canonical drivers have never been detected by any method, suggesting that they may elicit their role through non-mutational mechanisms or may fall below the detection limits of current approaches. Given the prevalence of cancer coding screens (**Figure 1A**), only coding driver alterations have been reported for most genes (**Figure 1E**) while 16% of them (531) were identified as drivers uniquely in noncoding screens. Since the prediction of drivers with noncoding alterations remains challenging, we further investigated the type of support that these genes had for their driver activity. The overwhelming majority of them (467 genes, 87%) have been predicted as drivers in only one screen. The remaining 64 genes are canonical drivers, have been predicted as drivers in multiple screens or have additional experimental support for their driver activity (**Figure 1F**)

Applying a similar approach (**Supplementary Figure 1A-C, Additional File 2**), we reviewed 18 sequencing screens of healthy or diseased (noncancer) tissues. They collectively reported 95 genes whose somatic alterations could drive non-malignant clone formation (healthy drivers). Interestingly, only eight of them were not cancer drivers (**Figure 1G, Supplementary Table 3, Additional File 1**), suggesting high overlap between genetic drivers of cancer and noncancer evolution. However, since many noncancer screens only re-sequenced cancer genes or applied methods developed for cancer genomics (**Supplementary Figure 1E, Additional File 2**), this overlap may be overestimated.

### The ability to capture cancer but not healthy driver heterogeneity increases with the donor sample size

To compare cancer and healthy drivers across and within tissues, we grouped the 122 cancer types and 12 noncancer tissues into 12 and seven organ systems, respectively (Methods).

Despite the high numbers of sequenced samples (**Supplementary Table 4, Additional File 1**) and detected drivers (**Figure 1**), several lines of evidence indicated that our knowledge of cancer drivers is still incomplete. First, we detected a strong positive correlation between cancer drivers and donors overall (**Figure 2A**) and in individual organ systems (**Supplementary Figure 2, Additional File 2**). This suggests that the current ability to identify new drivers depends on the number of samples included in the analysis. Second, candidates outnumbered canonical drivers in all organ systems except those with small sample size or low mutation rate such as paediatric cancers, where only the most recurrent canonical drivers could be identified (**Figure 2B**). Third, large donor cohorts enabled detection of a broader representation of canonical drivers than small cohorts (**Figure 2C**). For example, pooling thousands of samples together led to >60% of canonical drivers being detected in adult pancancer re-analyses. Therefore, the size of the cohort influences the level of completeness and heterogeneity of the cancer driver repertoire. This is not surprising since all current approaches act at the cohort level, searching for positively selected genes altered more frequently than expected (**Supplementary Figure 1D, Additional File 2**).

**Figure 2.**
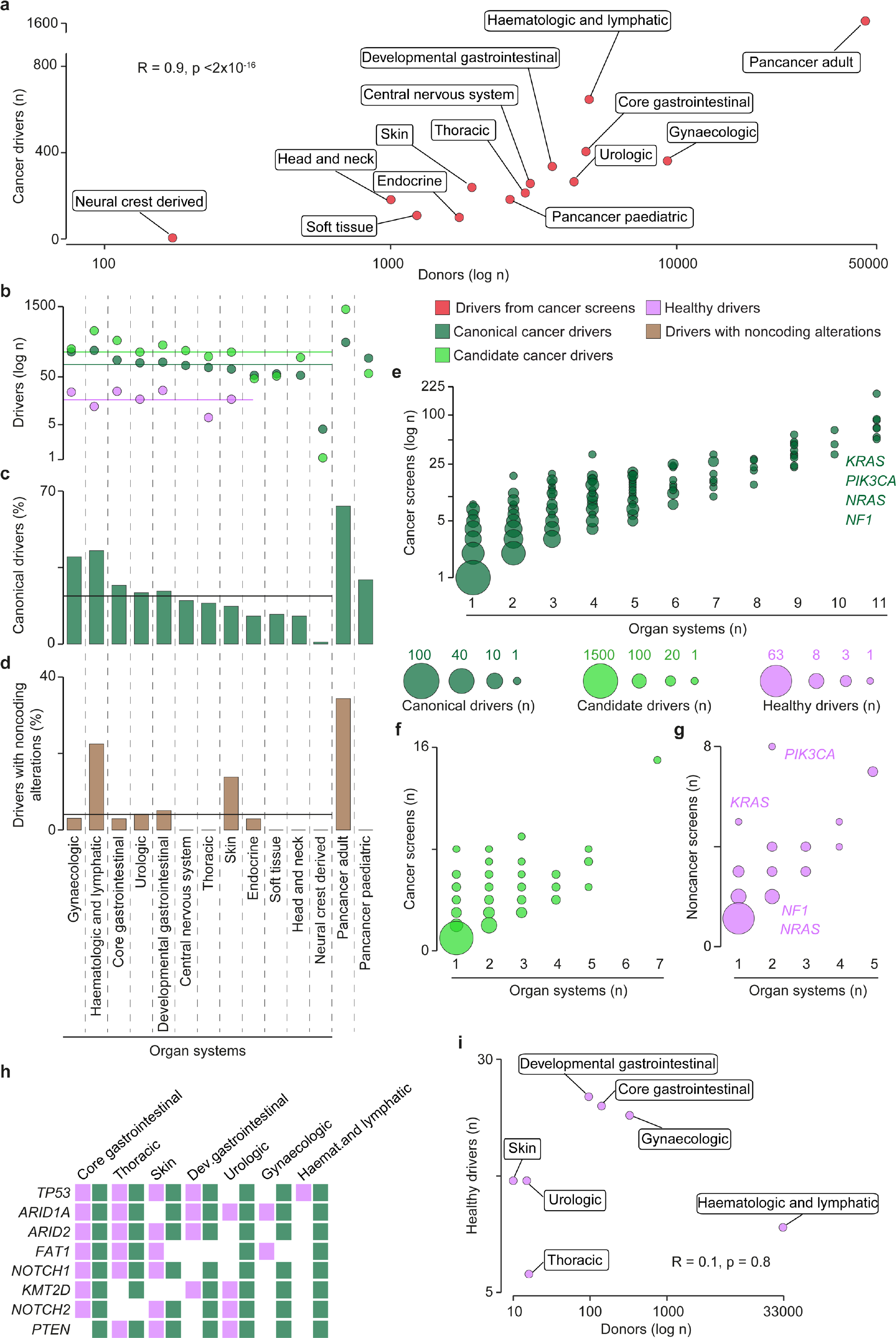
Distribution of driver annotations by organ system. **a**. Correlation between numbers of sequenced donors and identified cancer drivers across organ systems. Spearman correlation coefficient R and associated p-value are shown. **b**. Number of canonical, candidate and healthy drivers in each organ system. Horizontal lines indicate the median number of canonical (92), candidate (160) and healthy (17) drivers across organ systems. **c**. Proportion of canonical drivers detected in each organ system over canonical drivers detected in all cancer screens (421). The horizontal line indicates the median across all organ systems (22%). **d**. Proportion of genes with noncoding driver alterations over all cancer drivers in each organ system. The horizontal line indicates the median across all organ systems (4%). Number of canonical (**e**), candidate (**f**) and healthy (**g**) drivers across screens and organ systems. Representative genes with different recurrence between cancer and healthy tissues are indicated. **h**. Organ system distribution of the top eight recurrent healthy drivers. The full list is provided as **Supplementary Table 5, Additional File 1**. **i**. Correlation between numbers of sequenced donors and identified healthy drivers across organ systems. Spearman correlation coefficient R and associated p-value are shown.

Our analysis also showed that the contribution of noncoding driver alterations remains largely unappreciated and noncoding drivers have not yet been reported in several tumours, including all paediatric cancers (**Figure 2D**). Owing to the re-analysis of large whole genome collections ^21,22,23,24,25,26^, almost 40% of adult pancancer drivers were instead modified by noncoding alterations (**Figure 2D**). Haematological and skin tumours also had a high proportion of noncoding driver variants thanks to screens focused on noncoding mutations ^27,28^. Therefore, the re-analysis of already available whole genome data and further sequencing screens of noncoding variants are needed to fully appreciate their driver contribution.

Compared to cancer, sequencing screens of noncancer tissues are still in their infancy, as reflected by the lower numbers of screened tissues and detected drivers (**Figure 2B**). Despite this, some similarities and differences with cancer drivers could already be observed. Like cancer drivers (**Figure 2E-F, Supplementary Table 5, Additional File 1**), also healthy drivers were mostly organ-specific (**Figure 2G**) and the most recurrent healthy drivers were also cancer drivers in the same organ system (**Figure 2H, Supplementary Table 5, Additional File 1**). However, some recurrent cancer drivers (*KRAS, PI3KCA, NRAS, NF1*) were reported to drive noncancer clonal expansion only in one or two organ systems (**Figure 2G**). Therefore, differences start to emerge at the tissue level between drivers of cancer and noncancer evolution. Moreover, unlike cancer drivers, no correlation existed between numbers of drivers and donors (**Figure 2I**). This is likely affected by the lower number of noncancer sequencing studies available so far. If additional studies will confirm the absence of correlation, this may indicate that the healthy driver repertoire is easier to saturate since less drivers are needed to initiate and sustain noncancer clonal expansion ^10,11^.

### Alteration pattern hints at driver mode of action and confirms the incompleteness of the driver repertoire

To gain further insights into their mode of action, we mapped the type of alterations acquired by cancer and healthy drivers in 34 cancer types from TCGA. After predicting the damaging alterations in 7953 TCGA samples with matched mutation, copy number and gene expression data (Methods), we identified the drivers with loss-of-function (LoF) and gain-of-function (GoF) alterations in these samples, respectively (**Figure 3A**).

**Figure 3.**
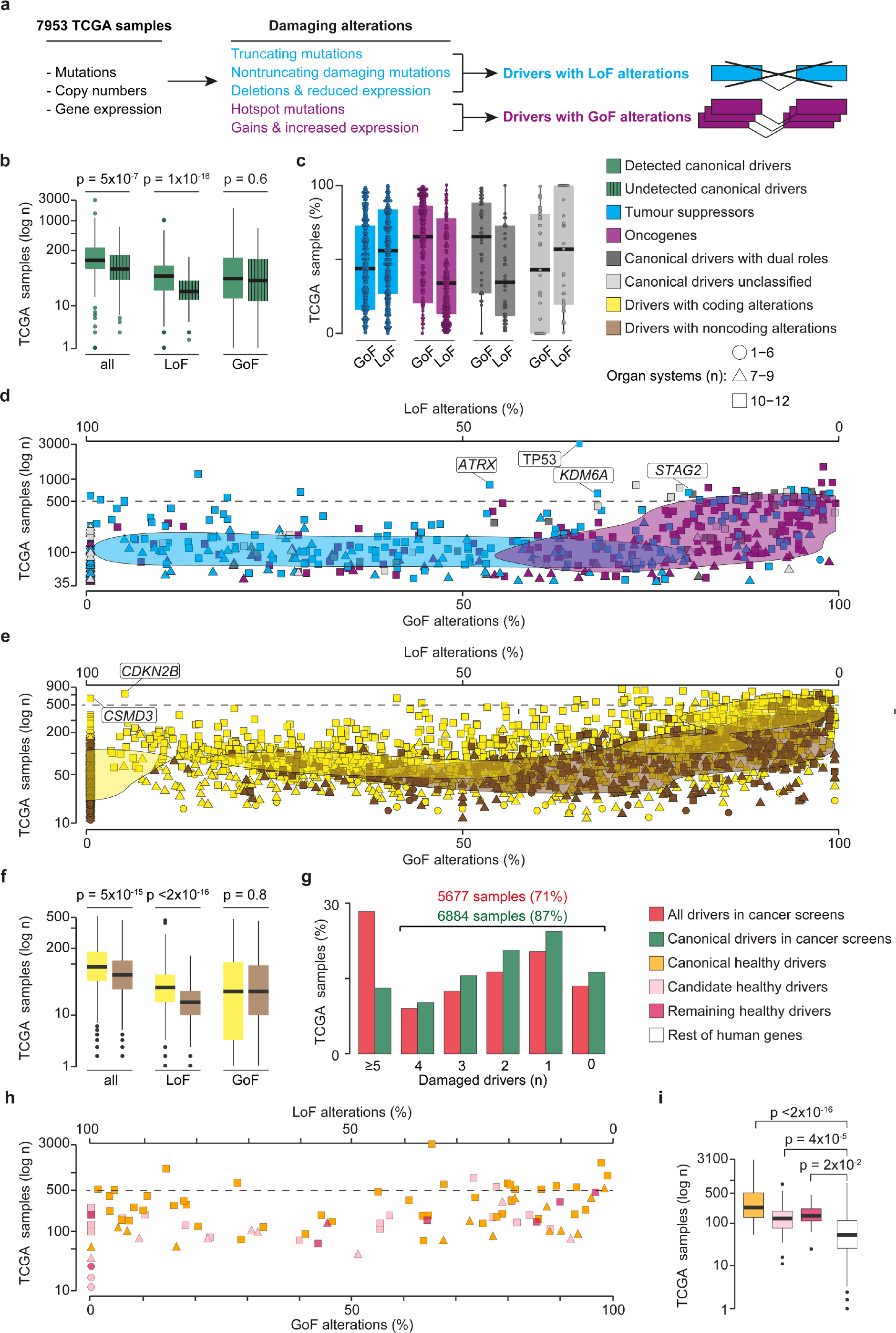
Damaging alteration pattern of drivers in TCGA. **a**. Identification of damaged drivers in 7953 TCGA samples. Mutations, gene deletions and amplifications were annotated according to their predicted damaging effect. This allowed to distinguish drivers acquiring loss-of-function (LoF) or gain-of-function (GoF) alterations. **b**. Number of TCGA samples with damaging alterations (all, LoF, GoF) in canonical drivers that were detected (421) or undetected (170) by cancer driver detection methods. **c**. Proportion of TCGA samples with GoF and LoF alterations in tumour suppressors, oncogenes and canonical drivers with a dual or unclassified role. Proportion of TCGA samples with GoF and LoF alterations in (**d**) canonical drivers and (**e**) **d**. candidate drivers. Genes mentioned in the text are highlighted. The two-dimensional gaussian kernel density estimations were calculated for each driver group using the R density function. **f**. Number of TCGA samples with damaging alterations (all, LoF, GoF) in drivers previously reported in coding and noncoding sequences. **g**. Proportion of samples with variable numbers of all damaged drivers or only canonical drivers. **h**. Proportion of TCGA samples with GoF and LoF alterations in healthy drivers. Canonical and candidate healthy drivers correspond to genes with a known or predicted cancer driver role. **i**. Number of TCGA samples with damaged canonical, candidate and remaining healthy drivers and the rest of human genes. All distributions were compared using a two-sided Wilcoxon rank-sum test.

The comparison between canonical cancer drivers detected and undetected in sequencing screens (**Figure 1D**) revealed that the latter were damaged in a significantly lower number of samples, due to fewer LoF alterations (**Figure 3B, Supplementary Figure 3A, Additional File 2**). GoF alterations were instead comparable between the two groups, suggesting that current driver detection methods fail to identify drivers that undergo copy number gains but are rarely mutated.

We confirmed that the driver alteration patterns reflected their mode of action, with canonical tumour suppressors and oncogenes showing a prevalence of LoF and GoF alterations, respectively (**Figure 3C**). Canonical drivers with a dual role resembled the alteration pattern of oncogenes while those still unclassified had a prevalence of LoF alterations, suggesting a putative tumour suppressor role (**Figure 3C**). While all frequently altered (>500 samples) oncogenes were overwhelmingly modified by GoF alterations (**Supplementary Table 6, Additional File 1**), 16 of the 22 most frequently altered tumour suppressors had a prevalence of GoF alterations (**Figure 3D**). In the majority of cases this was due to different alteration patterns across organ systems (**Supplementary Figure 3B, Additional File 2**) and a possible oncogenic role has been documented for some others ^29,30,31,32,33,34,35,36,37,38^.

Since candidate drivers had no annotation of their mode of action, we reasoned that their alteration pattern could hint at their role as tumour suppressors or oncogenes. According to their prevalent pancancer alterations, 1318 candidates could be classified as putative tumour suppressors and 1405 as putative oncogenes (**Supplementary Table 6, Additional File 1**). Interestingly, while candidates with predicted coding driver alterations showed similar distributions of LoF and GoF alterations (**Figure 3E**), those with only noncoding driver alterations had significantly lower occurrence of LoF alterations (**Figure 3F, Supplementary Figure 3C, Additional File 2**). This may suggest an activating role for their noncoding alterations too. Almost all candidates damaged in ≥500 samples (111/115) were putative oncogenes (**Figure 3E, Supplementary Table 6, Additional File 1**). Of the four putative tumour suppressors, *CSMD3* has a disputed cancer role ^39,40,41^ and a likely inflated mutation rate ^42^, while *CDKN2B* cooperates with its paralog *CDKN2A* to inhibit cell cycle ^43^, supporting its tumour suppressor role.

The number of damaged cancer drivers in individual TCGA samples confirmed that, despite all efforts, the current driver repertoire is still largely incomplete. The large majority of samples (71% and 87%, considering all drivers or only canonical drivers, respectively) had less than five damaged drivers and ∼15% of them had no damaged driver (**Figure 3G**).

Given their high overlap with cancer drivers, most healthy drivers were recurrently damaged in cancer samples with no prevalence of GoF or LoF alterations (**Figure 3H, Supplementary Table 6, Additional File 1**). Interestingly, all healthy drivers, even the eight with no cancer involvement, were damaged in significantly more cancer samples than the rest of human genes (**Figure 3I**). Moreover, 57% of TCGA samples had at least two altered drivers, one of which was a healthy driver, further supporting the hypothesis that more than one driver may be needed to promote transformation of nonmalignant clones into cancer ^10,11^.

### Properties of cancer and healthy drivers support their central role in the cell

A substantial body of work including our own ^44,45,46,47,48,49,50,51,52,53^ has shown that cancer drivers differ from the rest of genes for an array of systems-level properties (**Figure 1A**) that are consequence of their unique evolutionary path and role in the cell. Using our granular annotation of drivers, we set out to check for similarities and differences across driver groups.

We confirmed that cancer drivers, and in particular canonical drivers, were more conserved throughout evolution and less likely to retain gene duplicates than other human genes (**Figure 4A, Supplementary Table 7, Additional File 1**). They also showed broader tissue expression, engaged in a larger number of protein complexes, and occupied more central and highly connected positions in the protein-protein and miRNA-gene networks (**Figure 4A**). We reported substantial differences between tumour suppressors and oncogenes, with the former enriched in old and single-copy genes showing broader tissue expression (**Figure 4B, Supplementary Table 7, Additional File 1**).

**Figure 4.**
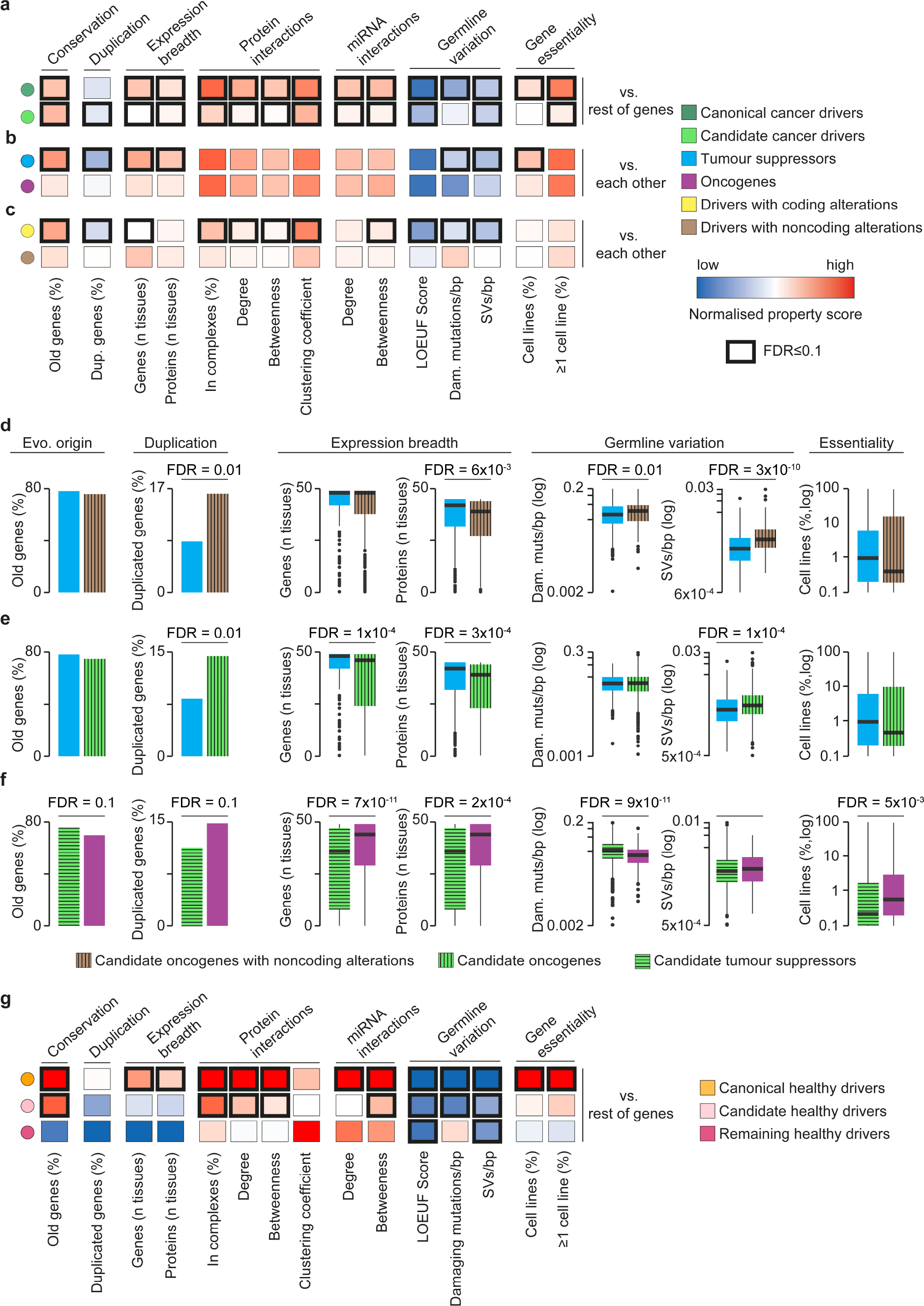
Systems-level properties of cancer and healthy drivers. Comparisons of systems-level properties between (**a**) canonical or candidate cancer drivers and the rest of human genes; (**b**) tumour suppressors and oncogenes, (**c**) cancer genes with coding driver alterations and cancer genes with noncoding driver alterations. The normalised property score was calculated as the normalised difference between the median (continuous properties) or proportion (categorical properties) values in each driver group and the rest of human genes (Methods). Comparisons of systems-level properties between (**d**) candidate oncogenes with noncoding driver alterations (324) and canonical tumour suppressors; (**e**) candidate oncogenes (1405) and canonical tumour suppressors; (**f**) candidate tumour suppressors (1318) and canonical oncogenes. **g**. Comparisons of systems-level properties between canonical healthy, candidate healthy and remaining healthy drivers and the rest of human genes. Proportions of old (pre-metazoan), duplicated, essential genes, and proteins involved in complexes were compared using a two-sided Fisher’s exact test. Distributions of gene and protein expression, protein-protein, miRNA-gene interactions, and germline variation were compared using a two-sided Wilcoxon rank-sum test. False discover rate (FDR) was corrected for using Benjamini-Hochberg.

We further expanded the systems-level properties of cancer drivers by exploring their tolerance towards germline variation, because this may indicate their essentiality. Using germline data from healthy individuals ^54^, we compared the loss-of-function observed/expected upper bound fraction (LOEUF) score, which quantifies selection towards LoF variation ^54^ as well as the number of damaging mutations and structural variants (SVs) per coding base pairs (bp) between drivers and the rest of genes (Methods). Cancer drivers, and in particular canonical drivers, had a significantly lower LOEUF score and retained fewer damaging germline mutations and SVs than the rest of genes (**Figure 4A**). This indicates that they are indispensable for cell survival in the germline. Selection against harmful variation was stronger in tumour suppressors than oncogenes (**Figure 4B**). This was supported by a significantly higher proportion of cell lines where cancer drivers, and in particular tumour suppressors, were essential (**Figure 4A**-B), as gathered from the integration of nine genome-wide essentiality screens ^55,56,57,58,59,60,61,62,63^ (Methods).

Genes with noncoding driver alterations had weaker systems-level properties than those with coding alterations (**Figure 4C, Supplementary Table 7, Additional File 1**) and the subset of them with >50% GoF alterations resembled the property profile of oncogenes when compared to tumour suppressors (**Figure 4D, Supplementary Table 7, Additional File 1**). In general, all candidate drivers with a prevalence of GoF were similar to oncogenes, showing higher proportion of duplicated genes, narrower tissue expression, and higher tolerance to germline variation than tumour suppressors (**Figure 4E, Supplementary Table 7, Additional File 1**). Conversely, candidate drivers with a prevalence of LoF were older, less duplicated and less tolerant to germline variation than oncogenes (**Figure 4F, Supplementary Table 7, Additional File 1**).

Systems-level properties of healthy drivers varied according to the overlap with cancer drivers (**Figure 4G, Supplementary Table 7, Additional File 1**). Intriguingly, canonical healthy drivers showed stronger systems-level properties than any other group of drivers. In particular, they were enriched in evolutionarily conserved and broadly expressed genes encoding highly inter-connected proteins are regulated by many miRNA. Moreover, these genes showed a strong selection against germline variation and high enrichment in essential genes (**Figure 4G**). They therefore represent a core of genes with a very central role in the cell, whose modifications are not tolerated in the germline but are selected for in somatic cells because they confer selective growth advantages. Candidate healthy drivers and those not involved in cancer had a substantially different property profile (**Figure 4G**). Although numbers are too low for any robust conclusion, it is tempting to speculate that genes able to initiate noncancer clonal expansion but not tumourigenesis may follow a different evolutionary path.

### The Network of Cancer Genes: an open-access repository of annotated drivers

We collected the whole repertoire of 3347 cancer and 95 healthy drivers, their literature support and properties in the seventh release of the Network of Cancer Genes and Healthy Drivers (NCG^HD^) database. NCG^HD^ is accessible through an open-access portal that enables interactive queries of drivers (**Figure 5A**) as well as the bulk download of the database content.

**Figure 5.**
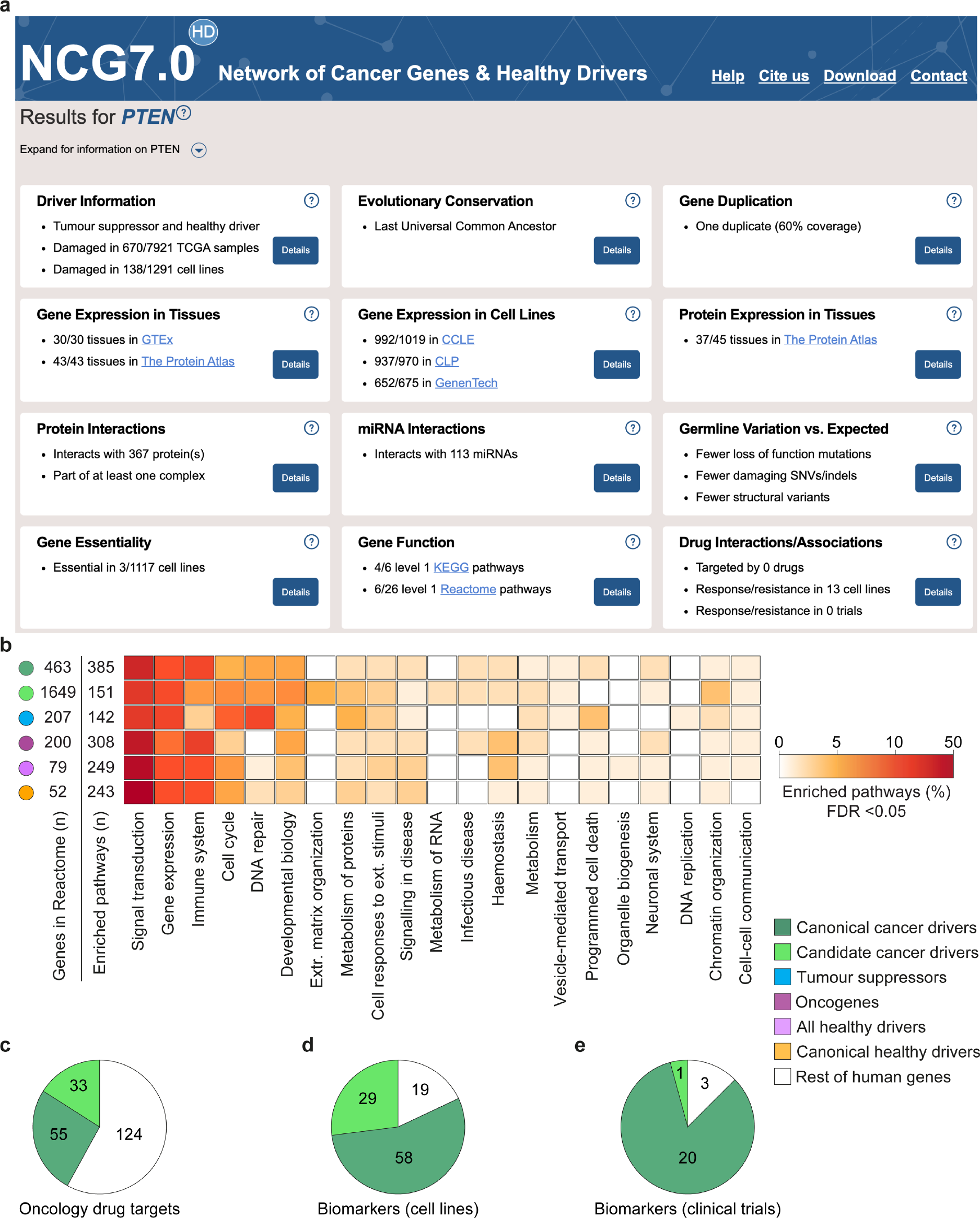
NCG^HD^ annotations of driver genes. **a**. Example of the type of annotation provided in NCG^HD^ for cancer and healthy drivers (in this case *PTEN*). Annotation boxes can be expanded for further details, with the possibility of intersecting data interactively (for example in the case of protein-protein or miRNA-gene interactions) and downloading data for local use. **b**. Proportion of Reactome levels 2-8 enriched pathways mapping to the respective level 1 in each driver group. Enrichment was measured comparing the proportion of drivers in each pathway to that of the rest of human genes with a one-sided Fisher’s exact test. FDR was calculated using Benjamini-Hochberg. The numbers of drivers and enriched Reactome pathways are reported for each group. Proportion of canonical and candidate cancer divers and rest of genes that are (**c**) targets of FDA approved antineoplastic drugs or biomarkers of response or resistance to oncological drugs in (**d**) cancer cell lines and (**e**) clinical studies. The corresponding numbers for each group are also shown.

In addition to the known or predicted mode of action and systems-level properties of cancer and healthy drivers, NCG^HD^ 7.0 also annotates their function, alteration pattern and gene expression profile in TCGA and cancer cell lines, reported interactions with antineoplastic drugs and potential role as treatment biomarkers (**Figure 5B**). Altogether this constitutes an extensive compendium of annotation of driver genes, including information relevant for planning experiments involving them.

Functional gene set enrichment analysis showed that at least 60% of enriched pathways (FDR <0.05) in any driver group converge to five broad functional processes (signal transduction, gene expression, immune system, cell cycle and DNA repair, **Figure 5B, Supplementary Table 8, Additional File 1**). Within these, tumour suppressors showed a prevalence in cell cycle and DNA repair pathways, while oncogenes were enriched in gene expression and immune system-related pathways (**Supplementary Table 8, Additional File 1**). Healthy drivers closely resembled the functional profile of cancer drivers, given the high overlap (**Figure 5B**). Because of the low number, it was not possible to assess the functional enrichment of healthy drivers not involved in cancer.

More than 9% of canonical cancer drivers are targets of anti-cancer drugs and cancer drivers constitute around 40% of their targets (**Figure 5C**). Moreover, most of the genes used as biomarkers of resistance or response to treatment in cell lines (**Figure 5D**) or clinical trials (**Figure 5E**) are cancer drivers, with an overwhelming prevalence of canonical cancer drivers.

## DISCUSSION

The wealth of cancer genomic data and the availability of increasingly sophisticated analytical approaches for their interpretation have substantially improved the understanding of how cancer starts and develops. However, our in-depth analysis of the vast repertoire of drivers that have been collected so far shows clear limits in the current knowledge of the driver landscape.

The identification of drivers as genes under positive selection or with a higher than expected mutation frequency within a cohort of patients has biased the current cancer driver repertoire towards genes whose coding point mutations or small indels frequently recur across patients. This strongly impairs the ability to map the full extent of driver heterogeneity leading to an underappreciation of the driver contribution of rarely altered genes and those modified through noncoding or gene copy number alterations, particularly amplifications. It also results in a sizeable fraction of samples with very few or no cancer drivers. This gap can be solved by complementing cohort-level approaches with methods that account for all types of alterations and predict drivers in individual samples, for example identifying their network deregulations ^64,65,66^ or applying machine learning to identify driver alterations ^67^. Alternatively, we have shown that systems-level properties capture the main features of cancer drivers, justifying their use for patient-level driver detection ^68,69^.

Our comprehensive study has also shown that cancer sequencing screens have so far mostly focused on resequencing and analysing the protein-coding portion of cancer genomes, leaving the contribution of noncoding drivers mostly uncovered. This bias may be addressed by performing additional cancer whole genome sequencing screens and improving analytical methods for the prediction of noncoding driver alterations.

Biases are starting to emerge also in the knowledge of healthy drivers. Many noncancer sequencing screens only targeted cancer genes and healthy driver detection methods used so far were originally developed for cancer genomics. Both these factors may contribute at least in part to explain the high overlap between drivers of cancer and noncancer evolution. An unbiased investigation of altered genes able to promote clonal expansion but not tumourigenesis could confirm whether their properties are indeed different from cancer drivers as suggested by our initial analysis on the few of them that have been identified so far. Additionally, the investigation of somatically mutated clones in noncancer tissues has just started and new screens are continuously published. The integrated analysis of these new studies will broaden our understanding of noncancer clonal expansion and further clarify its relationship with cancer transformation.

Our literature review did not cover driver genes deriving from chromosomal rearrangements or epigenetic changes because of their scattered annotations in the literature and difficulty in mapping their properties. Adding these genes to the repertoire when their knowledge will be mature will help closing the gaps in the knowledge of the genetic drivers of tumourigenesis.

## CONCLUSIONS

Our comprehensive analysis of cancer sequencing screens showed that the current repertoire of cancer driver genes is still incomplete and biased towards frequent mutations altering the gene coding sequence. This calls for the need of additional screens and methods to identify further coding and noncoding cancer drivers at single patient resolution. We confirmed the central role of cancer drivers within the cell, which is reflected in their evolutionary path and is shared by the majority of known healthy drivers. Further sequencing screens of healthy tissues are needed to clarify whether this is a feature of all genes whose mutations can driver noncancer clonal expansion or there is a group of healthy drivers that underwent a different evolutionary path.

## METHODS

### Literature curation

A literature search was carried out in PubMed, TCGA (https://www.cancer.gov/tcga) and ICGC (https://dcc.icgc.org/) to retrieve cancer screens published between 2018 and 2020 (**Supplementary Figure 1A, Additional File 2**). This resulted in 135 coding and 154 noncoding cancer screens. Of these, only 80 and 37 were retained after examining abstracts and full text, respectively. Criteria for removal were absence of driver genes or driver detection methods and the impossibility to map noncoding driver alterations to genes. The 37 new cancer screens were added to 273 publications previously curated by our team ^70^, totalling 310 publications (**Supplementary Table 1, Additional File 1**). A similar literature search retrieved 24 sequencing screens of noncancer tissues publications, 18 of which were retained after abstract and full-text examination (**Supplementary Figure 1A, Additional File 2; Supplementary Table 1, Additional File 1**). Each paper was reviewed independently by two experts and further discussed if annotations differed to extract the list of driver genes, the number of donors, the type of screen (whole genome, whole exome, target gene resequencing), the cancer or noncancer tissues and the driver detection method (**Supplementary Figure 1B, Additional File 2**).

Canonical cancer drivers were extracted from two publications ^17,18^ and the Cancer Gene Census ^71^ v.91. In the latter case, all Tiers 1 and 2 genes were retained, except those from genomic rearrangements leading to gene fusion (**Supplementary Figure 1B, Additional File 2**). Collected genes were further classified as tumour suppressor, oncogene or having a dual role according to the annotation in the majority of sources. Genes with conflicting or unavailable annotation were left unclassified.

Drivers from cancer screens and canonical sources underwent further filtering (**Supplementary Figure 1C, Additional File 2**). First, they were intersected with a list of 148 possible false positives ^18,42^. After manual check of the supporting evidence, two drivers were retained as canonical, five were considered as candidates, and 41 were removed (**Supplementary Table 2, Additional File 1**). The three resulting lists (canonical drivers, drivers from cancer screens and healthy drivers) were intersected to annotate canonical drivers in cancer screens, remaining drivers in cancer screens (candidate cancer drivers), canonical healthy drivers, candidate healthy drivers, and remaining healthy drivers (**Supplementary Figure 1C, Additional File 2; Supplementary Table 3, Additional File 1**).

Cancer types and noncancer tissues were mapped to organ systems using previous classification ^72^. Cancer types not included in this classification were mapped based on their histopathology (retinoblastoma to central nervous system; vascular and peripheral nervous system cancers to soft tissue; penile tumours to urologic system).

### Pancancer TCGA data

A dataset of 7953 TCGA samples with quality-controlled mutation (SNVs and indels), copy number and gene expression data in 34 cancer types was assembled from the Genomic Data Commons portal l ^73^ (https://portal.gdc.cancer.gov/). Mutations were annotated with ANNOVAR ^74^ (April 2018) and dbNSFP ^75^ v3.0 and only those identified as exonic or splicing were retained. Damaging mutations included (1) truncating (stopgain, stoploss, frameshift) mutations; (2) missense mutations predicted by at least seven out of 10 predictors (SIFT ^76^, PolyPhen-2 HDIV ^77^, PolyPhen-2 HVAR, MutationTaster ^78^, MutationAssessor ^79^, LRT ^80^, FATHMM ^81^, PhyloP ^82^, GERP++RS ^83^, and SiPhy ^84^); (3) splicing mutations predicted by at least one of two splicing-specific methods (ADA ^75^ and RF ^75^) and (4) hotspot mutations identified with OncodriveCLUST ^85^ v1.0.0.

Copy Number Variant (CNV) segments, sample ploidy and sample purity values were obtained from TCGA SNP arrays using ASCAT ^86^ v.2.5.2. Segments were intersected with the exonic coordinates of 19756 human genes in hg19 and genes were considered to have CNV if at least 25% of their transcribed length was covered by a CNV segment. RNA-Seq data were used to filter out false positive CNVs. Putative gene gains were defined as copy number (CN) >2 times sample ploidy and the levels of expression were compared between samples with and without each gene gain using a two-sided Wilcoxon rank-sum test and corrected for multiple testing using Benjamini-Hochberg. Only gene gains with false discover rate (FDR) <0.05 were retained. Homozygous gene losses had CN = 0 and Fragments Per Kilobase per Million (FPKM) values <1 over sample purity. Heterozygous gene losses had CN = 1 or CN = 0 but FPKM values >1 over sample purity. This resulted in 2192832 redundant genes damaged in 7921 TCGA samples.

In total, 518115 genes were considered to acquire LoF alterations because they underwent homozygous deletion or had truncating, missense damaging, splicing mutations, or double hits (CN = 1 and LoF damaging mutation), while 1674717 genes were considered to acquire GoF alterations because they had a hotspot mutation or underwent gene gain with increased expression (**Figure 3A**).

### Systems-level properties

Protein sequences from RefSeq ^87^ v.99 were aligned to hg38 using BLAT ^88^. Unique genomic loci were identified for 19756 genes based on gene coverage, span, score and identity ^89^. Genes sharing at least 60% of their protein sequence were considered as duplicates ^46^.

Evolutionary conservation was assessed for 18922 human genes using their orthologs in EggNOG ^90^ v.5.0. Genes were considered to have a pre-metazoan origin (and therefore conserved in evolution) if they had orthologs in prokaryotes, eukaryotes, or opisthokonts ^53^.

Gene expression for 19231 genes in 49 healthy tissues was derived from the union of Protein Atlas ^91^ v.19.3 and GTEx ^92^ v.8. Genes were considered to be expressed in a tissue if their expression value was ≥1 Transcript Per Million (TPM). Protein expression for 13229 proteins in 45 healthy tissues was derived from Protein Atlas ^91^ v.19.3 retaining the highest value when multiple expression values were available.

A total of 542397 nonredundant binary interactions between 17883 proteins were gathered from the integration of five sources (BioGRID ^93^ v.3.5.185, IntAct ^94^ v.4.2.14, DIP ^95^ (February 2018), HPRD ^96^ v.9 and Bioplex ^97^ v.3.0). Data on 9476 protein complexes involving 8504 proteins were derived from CORUM ^98^ v.3.0, HPRD ^96^ v.9 and Reactome ^99^ v.72. Experimentally supported interactions between 14747 genes and 1758 miRNAs were acquired from miRTarBase ^100^ v.8.0 and miRecords ^101^ v.4.0. Degree, betweenness and clustering coefficient were calculated for protein and miRNA networks using the igraph R package ^102^ v.1.2.6.

The loss-of-function observed/expected upper bound fraction (LOEUF) score for 18392 genes was obtained from gnomAD ^54^ v.2.1.1. Germline mutations (SNVs and indels) were obtained from the union of 2504 samples from the 1000 Genomes Project Phase 3 ^103^ v.5a and 125748 samples from gnomAD ^54^ v.2.1.1. Mutations were annotated with ANNOVAR ^74^ (October 2019) and 18812 genes were considered as damaged using the same definitions as for TCGA samples. A total of 32558 germline SVs for 14158 genes were derived using 15708 samples from gnomAD ^54^ v.2.1.1. The numbers of damaging mutations and SVs per base pairs (bp) were calculated for each gene.

Essentiality data for 19013 genes in 1122 cell lines were obtained integrating three RNAi knockdown and six CRISPR Cas9 knockout screens ^55,56,57,58,59,60,61,62,63^. Genes with CERES ^57^ or DEMETER ^63^ scores <-1 or Bayes score ^104^ >5 were considered as essential.

Proportions of duplicated, pre-metazoan, essential genes and proteins engaging in complexes were compared between gene groups using two-sided Fisher’s exact test. Distributions of tissues where genes or proteins were expressed, protein and miRNA network properties, LOEUF scores, damaging mutations and SVs per bp were compared between gene groups using a two-sided Wilcoxon test. Multiple comparisons within each property were corrected using Benjamini-Hochberg. For each systems-level property in each driver group (*d*) a normalised property score was calculated as:

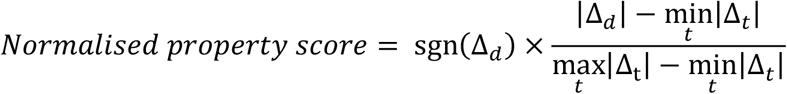

where *t* represents 11 gene groups (canonical drivers, candidate drivers, tumour suppressors, oncogenes, drivers with coding alterations, drivers with noncoding alterations, canonical healthy drivers, candidate healthy drivers, remaining healthy drivers and rest of human genes); sgn(Δ_*d*_) is the sign of the difference; and Δ_*d*_ indicates the difference of medians (continuous properties) or proportions (categorical properties) between each driver group and the rest of human genes. Minima and maxima were taken over all 11 gene groups for each property.

### Pancancer cell line data

Mutation, CNV and gene expression data for 1291 cell lines were obtained from DepMap ^56,105^ v. 20Q3. Mutations were functionally annotated using ANNOVAR ^74^ and LoF mutations were identified as described for TCGA samples. Hotspot mutations were detected using hotspot positions derived from TCGA. Homozygous gene deletions were defined as CN <0.25 times cell line ploidy and expression <1 TPM; heterozygous gene deletions were defined as 0.25< CN<0.75 times cell line ploidy; gene gains were defined as CN >2 times cell line ploidy and significantly higher expression relative to cell lines with no gene gains. Genes with LoF or GoF alterations were defined as for TCGA samples. To map cell lines to organ systems, they were first associated with the TCGA cancer types and then the same classification as for TCGA was used ^72^.

### Driver functional annotation

Gene functions were collected for 11778 proteins from Reactome ^99^ v.72 and KEGG ^106^v.94.1 (level 1 and 2). Driver enrichment in Reactome pathways (levels 2-8) compared to the rest of human genes was assessed using a one-sided Fisher’s exact test and corrected for multiple testing with Benjamini-Hochberg. Enriched pathways were then mapped to the corresponding Reactome level 1.

### Drug interactions

A total of 247 FDA approved, antineoplastic and immunomodulating drugs targeting 212 human genes were downloaded from DrugBank ^107^ v.5.1.8. Genetic biomarkers of response and resistance to drugs in cancer cell lines were obtained from Genomics of Drug Sensitivity in Cancer (GDSC) ^108^ v.8.2. Of those, only 467 associations with FDR ≤0.25 involving 129 drugs and 106 genes were retained. Genetic biomarkers of response and resistance in clinical studies were obtained from the Variant Interpretation for Cancer Consortium Meta-Knowledgebase ^109^ v.1. A total of 868 associations between drugs and genomic features involving 64 anti-cancer drugs and drug combinations and 24 human genes were retained ^109^.

### Database and website implementation

All annotations of driver genes were entered into a relational database based on MySQL ^110^ v.8.0.21 connected to a web interface enabling interactive retrieval of information through gene identifiers. The frontend was developed with PHP ^111^ v.7.4.15. The interactive displays of miRNA-gene and protein-protein interactions were implemented with the R packages Shiny ^112^ v.1.6.0 and igraph ^102^ v.1.2.6 and ran on Shiny Server v1.5.16.958.

## DECLARATIONS

### Ethics approval and consent to participate

Not applicable.

### Consent for publication

The results shown here are in part based upon data generated by the TCGA Research Network: https://www.cancer.gov/tcga.

### Availability of data and materials

The whole content of NCG^HD^ can be freely downloaded from the website (http://network-cancer-genes.org/). No license is required. All the custom scripts used for the study are available upon request.

Original data were obtained from the following online sources

1000 Genomes Project Phase 3 ^103^ v.5a:

https://www.internationalgenome.org/category/phase-3/

BioGRID ^93^ v.3.5.185: https://thebiogrid.org/

Bioplex ^97^ v.3.0: https://bioplex.hms.harvard.edu/interactions.php

CORUM ^98^ v.3.0: http://mips.helmholtz-muenchen.de/corum/}

Depmap ^59,60^ v20Q3: https://depmap.org/portal/

DIP ^95^ (February 2018): https://dip.doe-mbi.ucla.edu/dip/Main.cgi

DrugBank ^107^ v.5.1.8: https://go.drugbank.com/

EggNog ^90^ v.5: http://eggnog5.embl.de/#/app/home

GDSC ^108^ v.8.2: https://www.cancerrxgene.org/

GnomAD ^54^ v.2.1.1: https://gnomad.broadinstitute.org/

GTEx ^92^ v.8: https://gtexportal.org/home/

HPRD ^96^ v.9: https://www.hprd.org/

IntAct ^94^ v.4.2.14: https://www.ebi.ac.uk/intact/home

KEGG ^106^ v.94.1: https://www.genome.jp/kegg/

Meta-KB ^109^ v.1: https://cancervariants.org/

MiRecords ^101^ v.8.0: http://c1.accurascience.com/miRecords/

MiTarBase ^100^ v.4.0:

https://mirtarbase.cuhk.edu.cn/~miRTarBase/miRTarBase_2022/php/index.php

NCI Genomics Data Commons Portal ^73^: https://gdc.cancer.gov/

PICKLES ^61^: https://pickles.hart-lab.org/

Protein Atlas ^91^ v.19.3: https://www.proteinatlas.org/

Reactome ^99^ v.72: https://reactome.org/

RefSeq ^87^ v.99: https://www.ncbi.nlm.nih.gov/refseq/

### Author contributions

LD analysed protein-protein interactions, protein complex, gene essentiality and cancer cell line data. MB analysed gene conservation. MRK analysed gene duplicability. HM analysed TCGA data. MB and HM analysed miRNA-target interactions. GS analysed gene function, RNA and protein expression and drug interactions. AAS, LM, NW and DR curated the literature. JN analysed germline variation. GS, MB, JG and KA developed the database. MRK, HM, LD, MB, MP, PD and AS developed the website. LD, MB, MRK, HM, GS, AAS and FDC analysed the data. FDC conceived and supervised the study. MB, AAS, GS, and FDC wrote the manuscript with contributions from LD and HM. All authors reviewed and approved the manuscript.

## Acknowledgments

We thank Steve Hindmarsh and Stefan Boeing for their contribution to the development of the NCG database and website.

## ADDITIONAL FILES

**Additional file 1:** Supplementary tables (XLSX). Table S1. Publications describing driver genes; Table S2. Putative false positive cancer drivers; Table S3. Canonical cancer drivers; Table S4. Donors in cancer and noncancer sequencing screens; Table S5. Drivers reported in cancer and noncancer screens; Table S6. Cancer and noncancer drivers damaged in TCGA; Table S7. Systems-level properties of driver genes; Table S8. Proportion of enriched pathways across driver groups.

**Additional file 2:** Supplementary Figures (DOCX). Figure S1, Literature search, review and annotation workflow; Figure S2. Correlation between numbers of donors and cancer drivers in individual organ systems; Figure S3: Patterns of driver damaging alterations in TCGA samples.

