## Additional File 2 Supplementary Figures for "Comparative assessment of genes driving cancer and somatic evolution in noncancer tissues: an update of the NCG resource"

**Supplementary Figure 1:** Literature search, review and annotation workflow.

**Supplementary Figure 2:** Correlation between numbers of donors and cancer drivers in individual organ systems.

**Supplementary Figure 3:** Patterns of driver damaging alterations in TCGA samples.

**Supplementary table 1.** Publications describing driver genes

**Supplementary table 2.** Putative false positive cancer drivers

**Supplementary table 3:** List of canonical cancer drivers

**Supplementary Table 4:** Donors in cancer and noncancer sequencing screens

**Supplementary Table 5:** Drivers reported in cancer and noncancer screens

**Supplementary Table 6:** Cancer and noncancer drivers damaged in TCGA

**Supplementary Table 7:** Systems-level properties of driver genes

**Supplementary Table 8:** Proportion of enriched pathways across driver groups

**Supplementary Figure 1.** Literature search, review and annotation workflow.

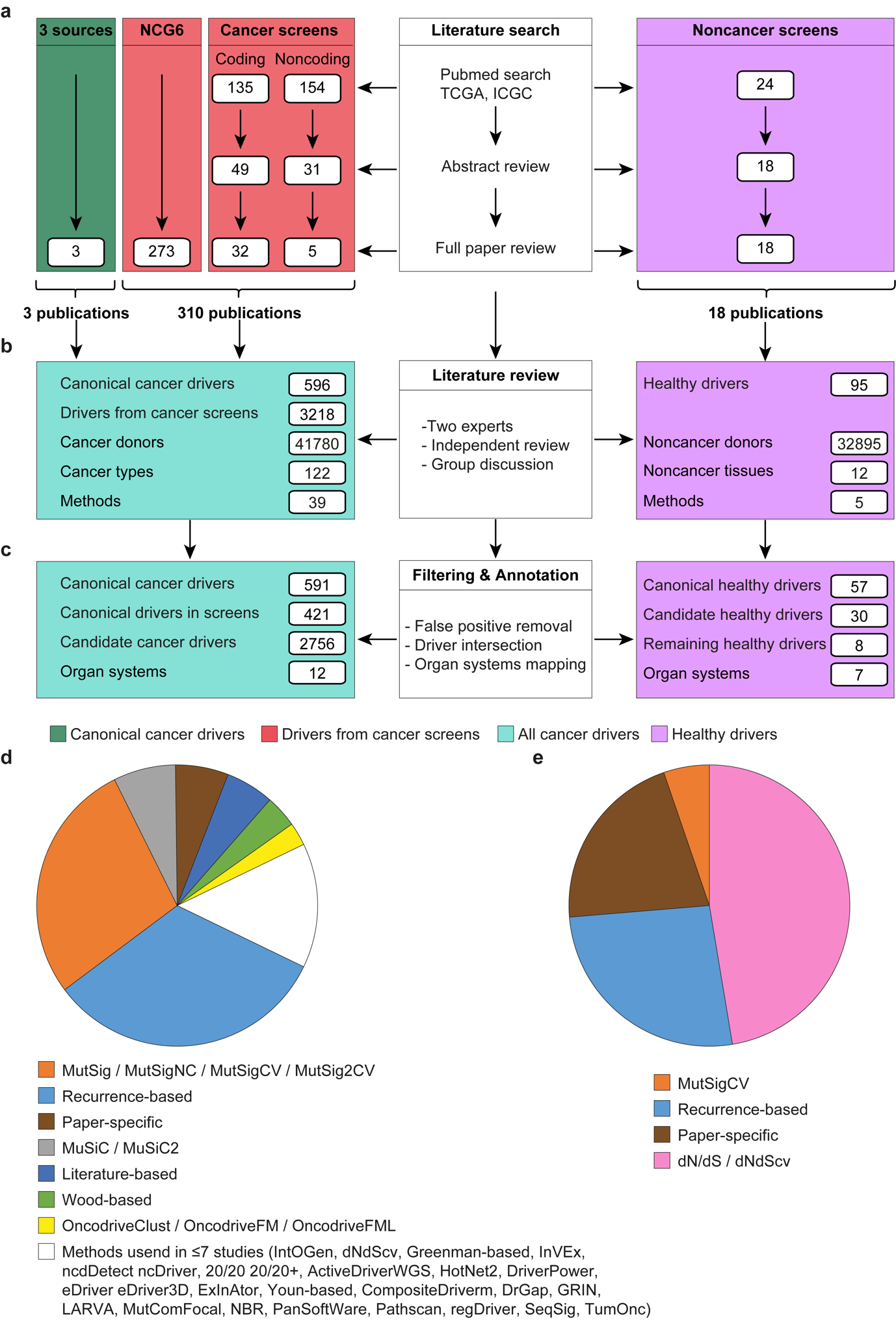

**a.** Publications reporting cancer and noncancer sequencing screens from the initial literature search in Pubmed, TCGA and ICGC were reviewed at the abstract and full text levels and added to three sources of canonical cancer genes and previously curated papers in NCG 6.0^1^ for a total of 331 publications.

**b.** Two experts reviewed each publication independently and conflicting annotations were further discussed. Lists of canonical cancer drivers, drivers from cancer and noncancer screens, cancer types and noncancer tissues, and methods used to detect drivers were annotated. Additionally, the number of cancer and noncancer donors were extracted.

**c.** The resulting lists of drivers were filtered out for possible false positives (Supplementary Table 2) and intersected to annotate the canonical drivers in cancer screens, candidate cancer drivers (remaining drivers in cancer screens), canonical healthy drivers, candidate healthy drivers, and remaining healthy drivers. Cancer types and noncancer tissues were mapped to organ systems^2^. The full workflow is explained in the Methods.

Usage of driver detection methods across (**d**) cancer and (**e**) noncancer screens. If a screen used several methods, it was counted multiple times. Multiple versions of the same method as well as methods used in less than 8 studies were aggregated. The full list is available in Supplementary Table 1.

**Supplementary Figure 2.** Correlation between numbers of donors and cancer drivers in individual organ systems

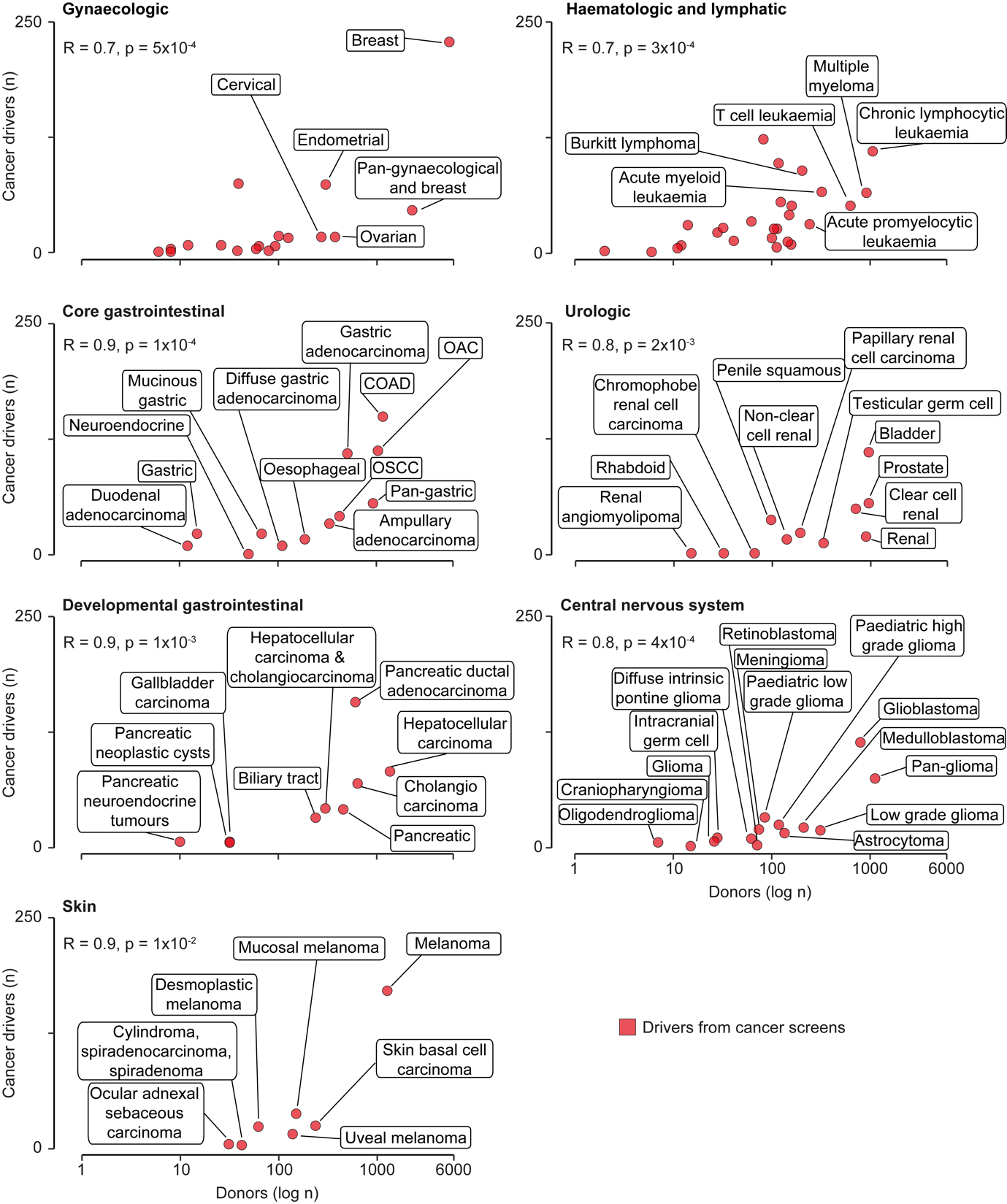

Correlations between numbers of sequenced donors and identified cancer drivers in individual cancer types mapping to each organ system. Only organ systems with significant correlations are reported. Spearman correlation coefficient R and associated p-value are shown. OAC: oesophageal adenocarcinoma. OSCC: oesophageal squamous cell carcinoma. COAD: colorectal adenocarcinoma.

**Supplementary Figure 3:** Patterns of driver damaging alterations in TCGA samples.

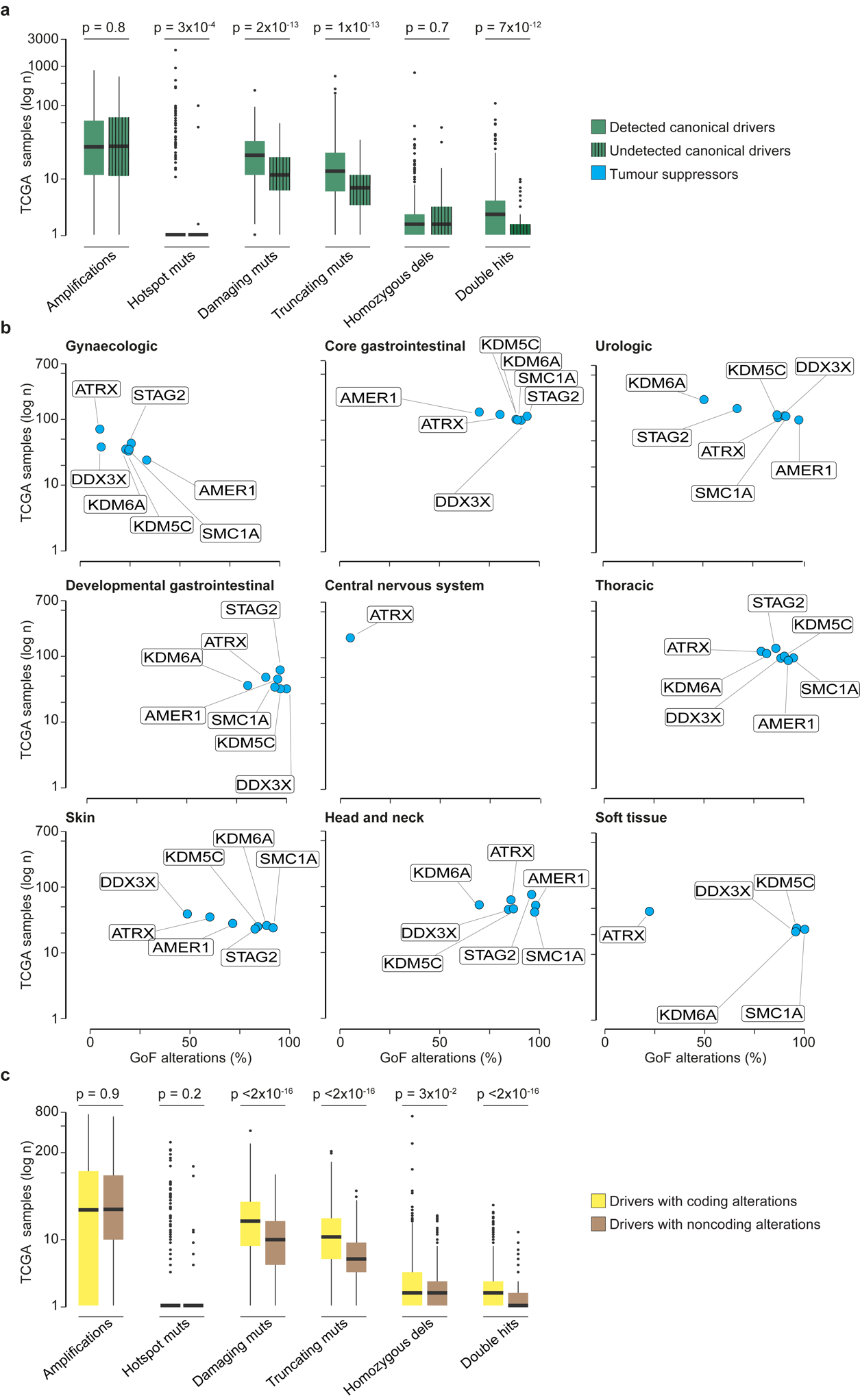

**a.** Number of TCGA samples with damaging alterations (all, LoF, GoF) in canonical drivers that were detected (421) or undetected (170) by cancer detection methods, divided by type of damaging alterations.

**b.** Proportion of gain of function (GoF) alterations affecting seven frequently damaged (>500 samples) canonical tumour suppressors. All these genes had an organ system- specific prevalence of GoF and loss of function (LoF) alterations.

**c.** Number of TCGA samples with damaging alterations in genes previously reported to have coding or only noncoding driver alterations, divided by type of damaging alterations.

All distributions were compared using a two-sided Wilcoxon rank-sum test.
